# Multi-scale planning helps resolve global conservation needs with regional priorities

**DOI:** 10.1101/2020.02.05.936047

**Authors:** D. Scott Rinnan, Yanina Sica, Ajay Ranipeta, John Wilshire, Walter Jetz

## Abstract

Area-based conservation through reserves or other measures is vital for preserving biodiversity and its functions for future generations^1–5^, but its effective implementation suffers from a lack of both spatial detail necessary for management practices^6^ and transparency around national responsibilities that might underpin cross-national support mechanisms^7^. Here we implement a conservation prioritization^2,8^ framework that accounts for spatial data limitations yet offers actionable guidance at a 1km resolution. Our multi-scale linear optimization approach delineates globally the areas required to meet area-based conservation targets for all ~32 000 described terrestrial vertebrate species, while offering flexibility in decision management to meet different local conservation objectives. Roughly 30.4% of land is sufficient to meet conservation targets for all species, of which 60.1% is either already protected^9^ or has minimal human modification^10^. However, the remaining 39.9% of human-modified areas need to be managed or restored in some form to ensure the long-term survival for over half of species. This burden of area-based conservation is distributed very unevenly among countries, and, without a process that explicitly addresses geopolitical inequity, meeting species conservation targets would require disproportionately large commitments from poorer countries (i.e., lower GNI). Our analysis provides baseline information for a potential intergovernmental and stakeholder contribution mechanism in service of a globally shared goal of sustaining biodiversity. Future updates and extensions to this global priority map have the potential to guide local and national advocacy and actions with a data-driven approach to support global conservation outcomes.

## Main Text

The current extinction crisis threatens biodiversity worldwide, driven primarily by loss of habitat due to human land use^5,11,12^. Negotiations are underway for a post-2020 Global Biodiversity Framework that provides an improved set of biodiversity targets for the coming decade and beyond^4,6,13,14^. Key principles shaping the new framework include a grounding in scientific understanding of the status and trends of the planet’s biodiversity, a focus on meaningful and measurable biodiversity outcomes, and development of mechanisms that support equitable management between parties^15^. Conservation policy and advocacy frequently features areal percentage targets (such 30% by 2030^15^) that have commonly been interpreted at the national or regional level, but generally fail to account for the uneven distribution of global biodiversity^6,16^. Other current CBD goals, by contrast, emphasize minimizing species extinctions and supporting global biodiversity persistence by enhancing perceptions of biodiversity importance. Thus, a good starting point for addressing issues of global conservation is one that reflects the multifaceted nature of these goals: given a shared objective to protect our planet’s biodiversity, what and where are the baseline amounts of area required, and what actions are needed to achieve it?

Several recent studies have addressed these questions by using species expert range maps to inform global conservation priorities^17–19^, and have featured increasingly finer degrees of spatial resolution of the type needed for translating globally-informed results into regional contexts and strategies^20^. Unfortunately, this pursuit of high-resolution maps often neglects or trivializes the spatial accuracy of the underlying species data; range maps interpreted at grain sizes less than 50 km inevitably suffer from geographically- and ecologically-variable false presences^21,22^. This can affect spatial prioritization approaches by preventing an unbiased and accurate quantification of each species’ area of occupancy and reserve coverage, and by overstating the precision with which high priority conservation locations can be identified^23^.

Here we introduce and apply a hierarchical prioritization framework that identifies areas for biodiversity conservation and leverages the spatial uncertainty of biodiversity data to help bridge the gap between global conservation objectives and local management practices. Our approach uses linear optimization^2,8^ (see Methods) to allocate sufficient habitat^24^ for all terrestrial vertebrate species, accounting for currently protected areas (PAs)^9^. For each species, we optimize for two different categories of habitat at two different spatial resolutions: general habitat via coarsely gridded expert range maps (ER), and finer scale habitat-suitable range (HSR) via expert maps refined by known habitat associations^12^. This approach provides more comprehensive species protection than using either habitat category alone: species with less HSR than ER, for example, have typically experienced some amount of habitat loss from human activity; these species will benefit from both preservation of current habitat (via HSR) and restoration of degraded habitat (i.e., ER that is not presently suitable). The solution recommends the optimal proportion of area to protect within each coarse-grain planning unit (PU; of uniform 770km^2^ size) without immediately resolving or prescribing the fine-scale locations within. Local management decisions can then be applied between and within different planning units, additionally informed by more detailed local data on populations and priorities as available, which in turn collectively determine the total amount of species habitat protected (Fig. 1D).

**Figure 1:**
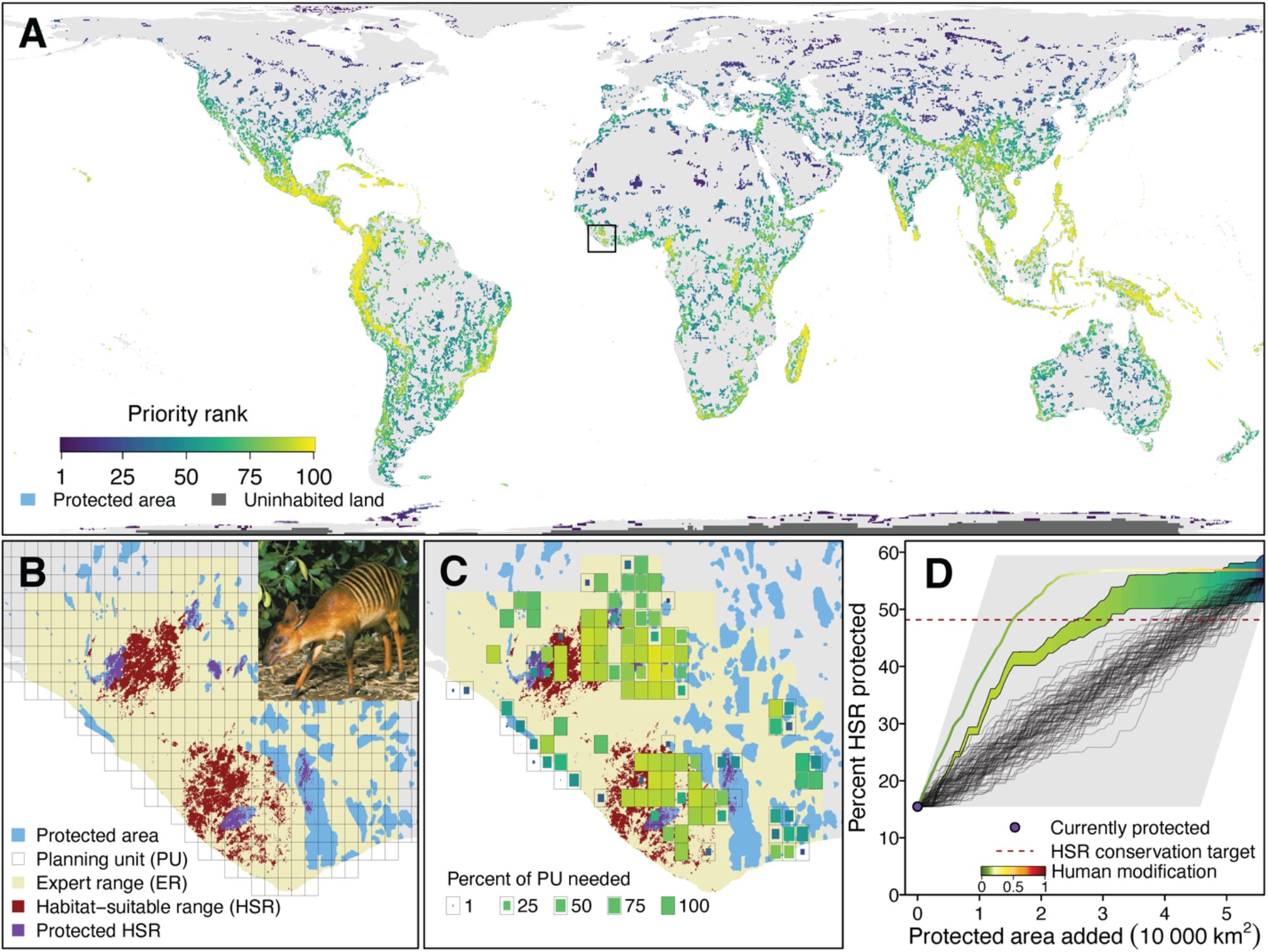
Conservation priorities for terrestrial vertebrates (32 649 species) and their supporting evidence and potential action at local scale. In (A) conservation priority areas are colored by the percentile rank of total network-size endemism, i.e., the proportion of the protected range of a species that is found in each planning unit (summed across all species). Displayed PU size reflects the proportion of land needed within each PU as determined by optimization. Dark grey regions indicate terrestrial PUs not overlapping with ranges of any study species. (B)-(D) illustrate the cross-scale planning implications for the zebra duiker (*Cephalophus zebra*), a threatened West Africa mammal species (see inset in A). (B) shows the Expert Range (ER) and habitat-suitable range (HSR) across PUs, with ~15% of each area currently conserved. (C) additionally displays the all-vertebrate priorities and ranks as identified in (A) that support fine-scale decision-making to address both this species and vertebrates at large. (D) Variability in conservation trajectories for zebra duiker HSR, depending on local conservation strategies. All trajectories will fall within the grey region. The widening colored band indicates the range of trajectories prioritizing PUs of higher priority rank first; within-PU selection of 1-km pixels will further determine trajectory path and total HSR protected, but all paths meet and exceed the 48% HSR conservation target. Black lines indicate trajectories determined by random selection of PUs and then within-PU pixels. The colored line indicates trajectory prioritizing 1-km pixels of lower human modification first, leading to more rapid conservation gains for the zebra duiker. Total area protected amounts to 40% of expert range and 56-60% of HSR, depending on choice of within-PU locations.

We identified 30.4% of global inhabited terrestrial surface area as needed to meet area-based conservation targets for 32 649 terrestrial vertebrate species, including the 14.2% of inhabited land that is already protected (Fig. 1A). Optimization tended to prioritize locations of higher species rarity^25^ and endemism^26^ (Fig. S1). 62.3% of PUs that were in the 90^th^ percentile of either endemism or rarity were selected for some amount of additional conservation, compared to just 17.6% of PUs in lower percentiles. To identify regions of higher conservation priority, we calculated the priority rank of each PU in the conservation area network as the proportion of the protected range of a species found within the PU, summed across all species. Priority rankings (Fig. 1A) generally reflected global patterns of terrestrial vertebrate endemism (r_s_ = 0.846, n = 42 485 for ER; r_s_ = 0.856, n = 40 881 for HSR), and their inter-taxonomic variation (Fig. S2) provides the taxon- and ultimately species-specific conservation hotspots that can help support advocacy or guide implementation at local levels^27^.

These PUs are further resolved to a finer spatial scale by selecting the specific 1-km pixels within each PU to meet the PU’s conservation area needs that collectively meet or exceed individual species targets globally. The selection process can reflect local management issues and concerns such as connectivity, expansion of current PAs, habitat integrity, political desirability, economic feasibility, carbon sequestration, and greater protection of high-profile species^28^. Figs. 1B-D illustrate this process for an individual example species and compare the trajectories of various local conservation strategies with 100 trajectories made by random selection of habitat. While all trajectories exceed both ER and HSR representation targets, those that target areas of higher priority rank or lower human modification^10^ (HM) tend to offer more rapid and overall habitat gains for species.

Fully resolving the global network by application of a uniform selection criterion yields further insights at higher resolution; minimizing the amount of within-network HM, for example, highlights the relevance of habitat quality (Fig. 2). Even as least modified areas are sought out first, only 4.7% of global areas selected for additional protection have no HM, and 11.6% have low, 26.6% moderate, 24.2% high, and 22.5% very high HM, respectively. This illustrates that comprehensive biodiversity conservation cannot be achieved simply by protecting areas of lower HM alone, but will require additional restoration efforts^10^. The degree of moderate- to very high-HM land identified as needing conservation also highlights the importance of other effective area-based conservation measures (OECMs)^3,29^, conservation in working lands, and efforts that include people^30,31^ as essential approaches to protect biodiversity.

**Figure 2.**
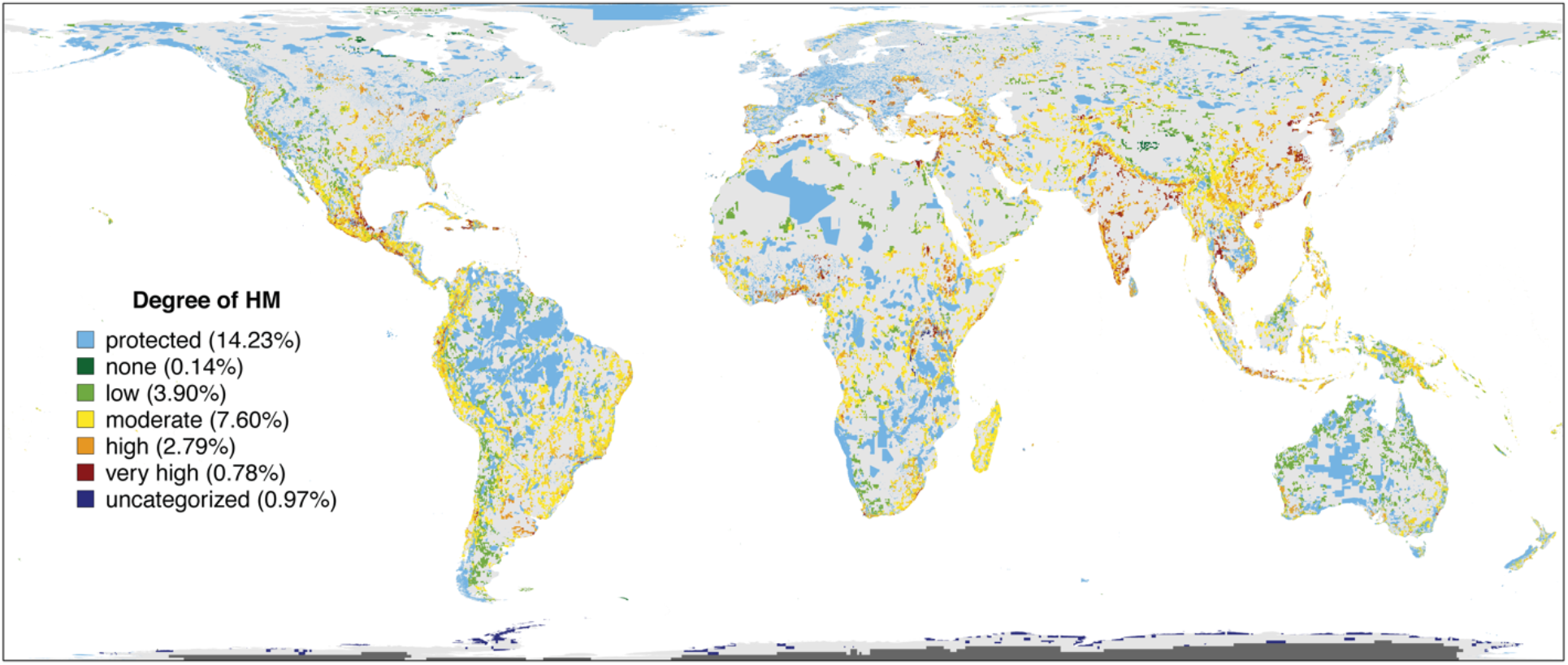
A putative 1-km conservation area network that provides protection for all terrestrial vertebrates while minimizing area needed and the degree of human modification (HM) contained therein. Amounts in parentheses indicate the total percentage of inhabited terrestrial surface area contained in the network by degree of HM; dark grey PUs without species occurrences were excluded from calculations of global area coverage. <1% of areas selected were not associated with an HM category, typified by regions of freshwater and ice.

Existing PAs meet ER conservation targets of only 27.1% of species, and HSR targets of only 22.6% of species (Fig. 3A, B). The minimum-HM approach suggests that adding areas of no and low HM (0–0.1) improves this to 40.9% of ER and 44.3% of HSR targets, and further adding areas of moderate HM (>0.1–0.4) meets 85.4% of ER and 94.9% HSR targets. Excluding areas of high HM (>0.4) fails to meet 15.1% of ER targets and of 5.1% of HSR targets (Fig. 3C, D), but half of these gap species are within 23.2% and 27.8% of their conservation targets, respectively. Although current PAs appear to safeguard less HSR than ER across taxa, rapid gains in HSR are achieved by the addition areas of low and moderate HM. This suggests diminishing returns on protecting areas of very high HM; indeed, since so little HSR exists within areas of very high HM, ER overlap with these areas may be an artifact of the spatial accuracy of the data.

**Figure 3.**
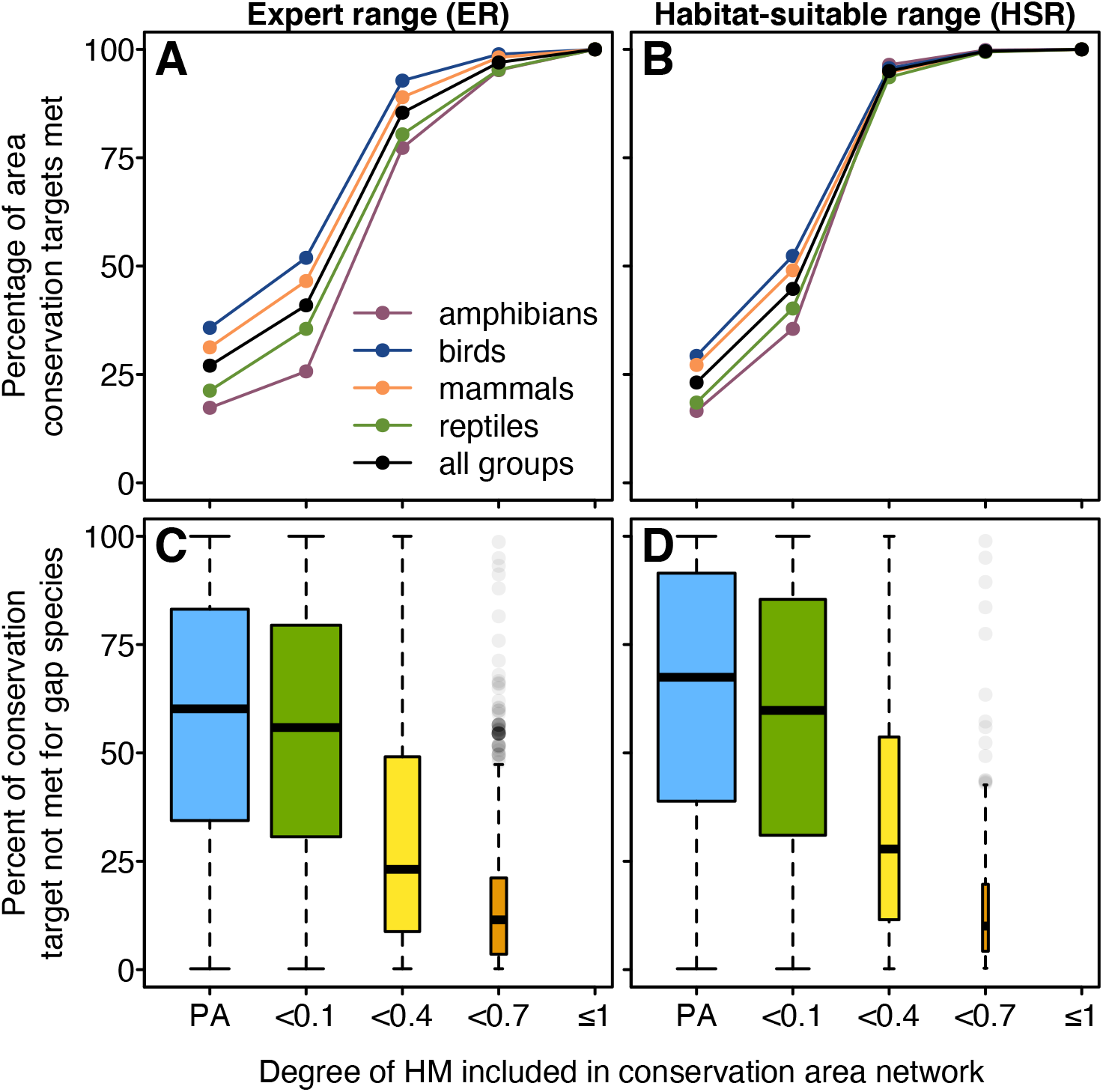
Variation in species coverage with the addition of increasingly modified areas. (A) The percentage of ER representation targets achieved by taxon when the network was constrained by differing degrees of HM. PA = currently protected areas only; <0.1 = PAs and areas with HM less than 0.1 (low); <0.4 = PAs and HM less than 0.4 (moderate); <0.7 = PAs and HM less than 0.7 (high); ≤1 = PAs and HM up to 1 (very high). (B) The percentage of HSR representation targets achieved by taxon. (C) Percent ER and (D) HSR representation targets that remain to be met for gap species. Boxplot width is proportional to the number of species represented in each HM category. Boxplot center lines indicate the medians, box limits the quartiles, whiskers 1.5x the interquartile ranges, and points the outliers.

Both relative and total areas required across HM categories differed strongly between ecoregions (Fig. 4A) and biomes (Fig. 4B). Percent of ecoregion needed and ecoregion area correlated negatively (r_s_ = −0.64, n = 847), and so did percent of biome needed and biome area (r_s_ = −0.92, n = 15), with some of the smallest biomes requiring the greatest relative area for conservation management. This underscores the limitations and inefficiency of any conservation targets — such as 17%, 30% or 50% — that are uniform across ecoregions, as commonly inferred from the expiring Aichi Biodiversity Target 11 or others more recently proposed^14^. The degree of HM in the network also varied greatly, further emphasizing the heterogeneity of the extent and distribution of restoration efforts necessary to ameliorate habitat degradation.

**Figure 4.**
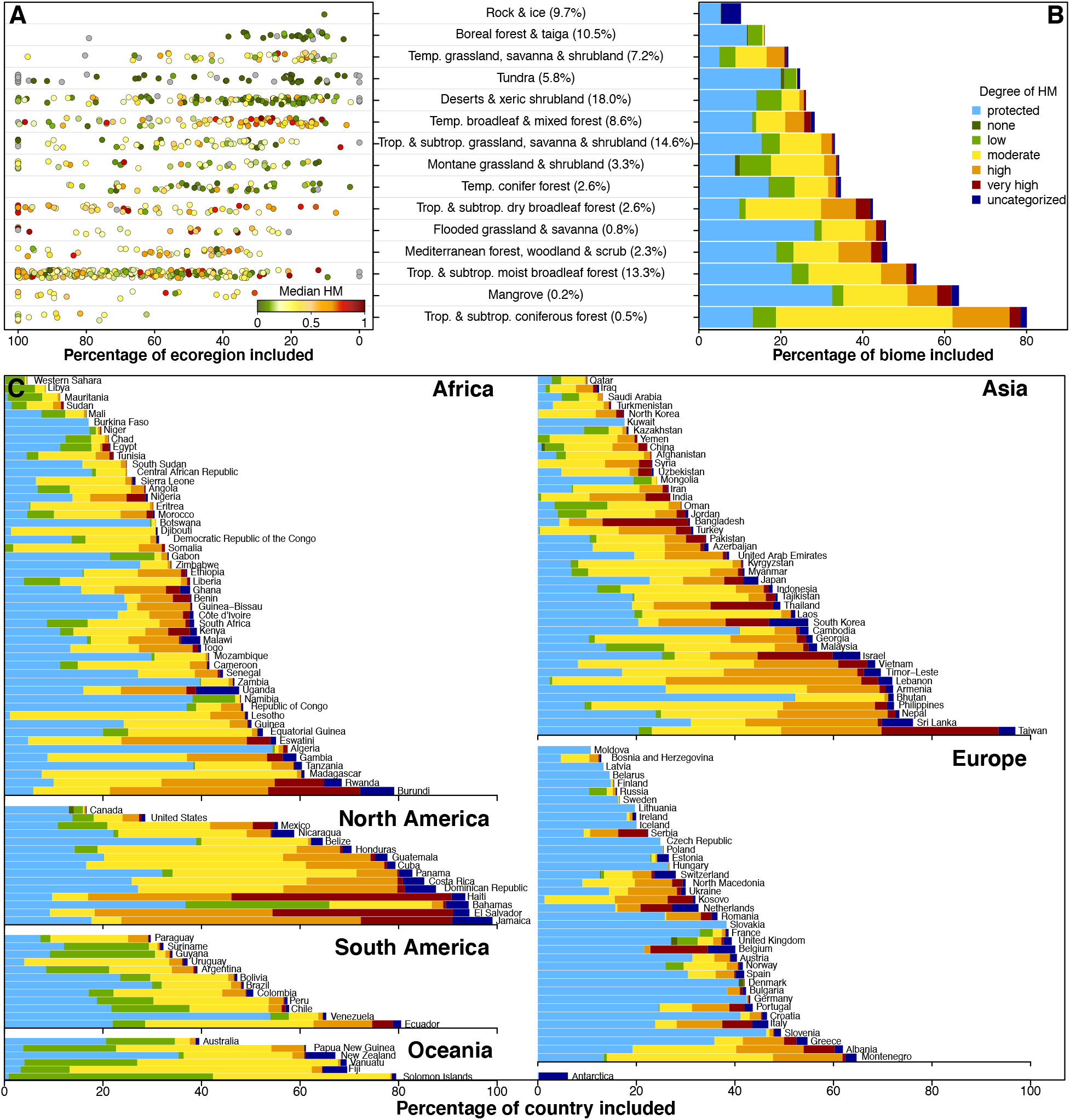
Conservation area network composition across ecoregions, biomes, and countries. (A) The percentage of each ecoregion contained in the network by biome category. Point colors indicate the median amount of HM in the network within each ecoregion. Percentages in parentheses indicate the total amount of global land area within each biome. (B) The percentage of each biome contained in the network by degree of HM and the amount currently protected. (C) Percentage of each country needed by degree of HM and current PAs.

Most relevantly, these considerable variances extend to political units: we found substantial disparities among countries and distinct administrative regions in the percent of land needed to meet conservation goals (Fig. 4C). Required commitments had a moderately negative correlation with country area (r_s_ = −0.36, n = 255), but were high for very small (e.g. island) nations. There was a similar relationship between countries in the amount of current HM in such a future conservation network, with a moderately negative correlation between area needed and mean HM (r_s_ = −0.37, n = 255). Generally, larger countries tended to have more low-HM land available for conservation.

Differences between nations were further amplified by the economic feasibility of achieving conservation objectives (Fig. 5). We characterized a country’s conservation burden as the area of its required conservation area network divided by its gross national income adjusted for purchasing power parity (GNI PPP), with total network area weighted by HM category; higher HM was weighted more heavily to reflect the costs associated with restoration of degraded habitat^32^. The most extreme national burdens were more than a hundred-fold larger or smaller than the global burden (i.e., global weighted-area needed over global GNI PPP). Countries with relatively little monetary wealth, such as Papua New Guinea and the Democratic Republic of the Congo, for example, require large percentages of land, and this problem is further exacerbated by the disproportionately high amount of degraded habitat that is needed to meet these goals. By contrast, many European countries require substantially smaller percentages of land, while having considerable financial resources to commit toward meeting conservation targets. Given that such disparities are often in part a direct result of a historical legacy and/or ongoing telecoupling of resource exploitation^33^, this underscores a globally-shared ethical responsibility toward addressing these inequities. Recognizing the preservation of species as a globally shared goal illustrates how the principle of “Common But Differentiated Responsibilities” from the United Nations Framework Convention on Climate Change could be applied to national targets for conservation in a framework that includes international support mechanisms^34^.

**Figure 5:**
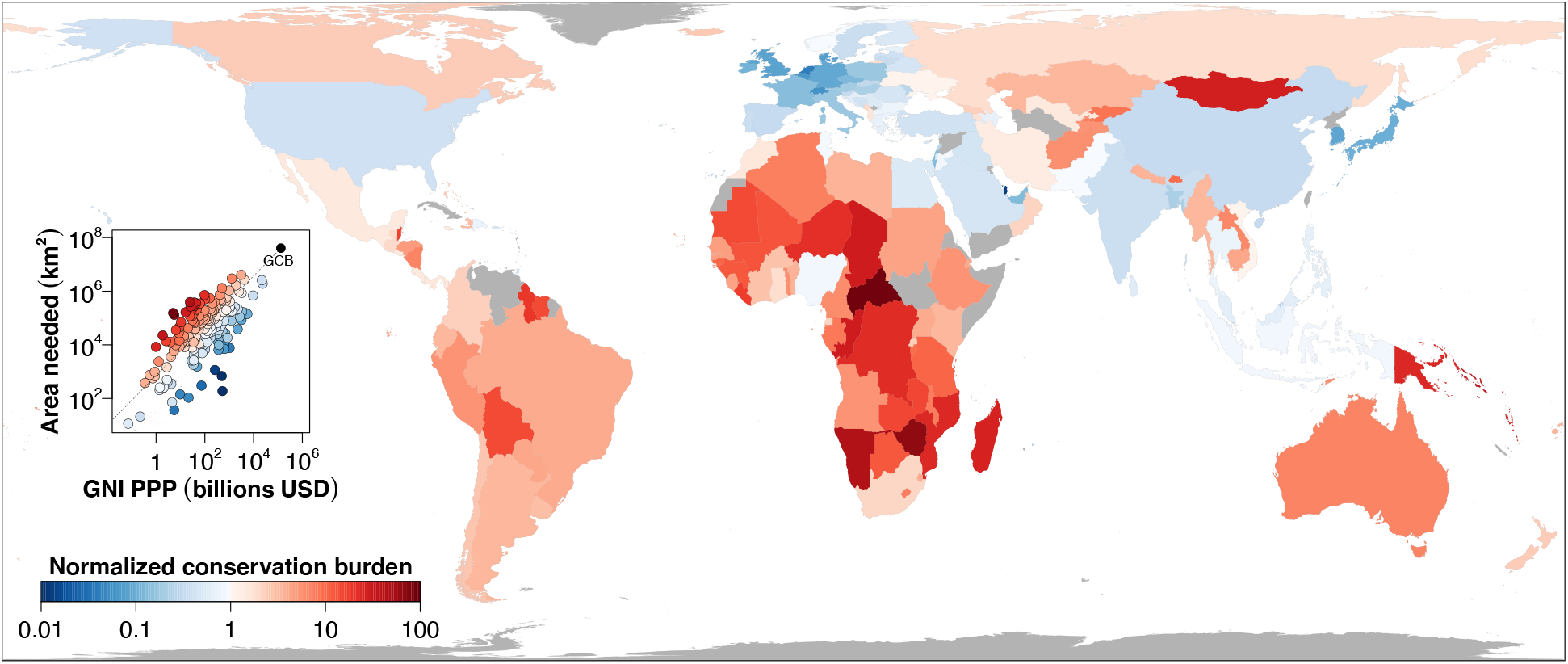
Differences in conservation burden between countries, calculated as a ratio of a weighted sum of each country’s area contained in the network and its gross national income adjusted for purchasing power parity (GNI PPP). The area sum is weighted by HM value. Color indicates national burdens normalized by the present global conservation burden (GCB) — representing the weighted sum of the entire network area divided by the sum of GNI PPP — which is equal to 0.367 m^2^/$. Countries in red have a higher burden than GCB, and countries in blue have a lower burden than GCB. Countries in grey did not have recent estimates of GNI PPP available.

Any successful and sustained conservation of the planet’s biodiversity necessitates a cooperative, coordinated international effort. The discussions of the CBD’s post-2020 Global Biodiversity Framework feature voluntary stakeholder contributions by Parties to the Convention and others as a vital mechanism for its implementation^15^. Our metric of conservation burden closely resembles the socioecological approach to intergovernmental biodiversity financing proposed by Droste *et al*.^7^, who found that a framework that accounted for the Human Development Index of countries best rewarded and incentivized global conservation action. In this light, conservation burden can be used as an implementation support mechanism of CBD goals by helping to guide intergovernmental fiscal subsidy agreements that encourage global conservation in the places it is most needed, while ensuring that economic and regulatory incentives promote biodiversity.

Our results are constrained by the still-limited knowledge of global biodiversity distributions^21,35,36^, requiring a tradeoff between taxonomic coverage and spatial resolution reflected in our analyses. Invertebrate and plant groups are vital for ecosystem health and function^37^ yet remain only partially mapped or are mapped over much coarser spatial units^38,39^. This prohibits their inclusion here, as any spatial differences in accuracy or representation would directly and non-transparently bias the resulting network. Previously identified similarities in endemism^26,40^ suggest that our proposed conservation area network would also help protect rare plant species. But such cross-taxon congruence is known to decrease rapidly toward finer spatial grains^41,42^, and both absolute and relative conservation burdens are expected to see slight shifts with the inclusion of other taxa. Efforts underway to develop the essential biodiversity information for species distributions for more groups and finer spatial grains through modeling and iterative feedback are poised to offer this vital support for outcome-focused global conservation^43^.

The presented baseline results and future updates provide a spatial blueprint for policy targets (such as those currently under renegotiation in the CBD) and initiatives (such as 30×30 and Half-Earth^44,45^) with stated goals of the preservation of species and their ecological diversity for future generations. Although area-based conservation measures may fall short of directly addressing drivers of biodiversity loss such as pollution and climate change^46^, protecting land from habitat loss and fragmentation remains one of the most effective means of conservation^1^. The priority places highlighted here ensure baseline goals of adequate global biodiversity representation but can be readily combined with other priority areas that address additional considerations such as wilderness^47,48^, carbon sequestration^19,49^, land-use change^48,50^, migration corridors^51^, and climatic refugia^52^, and can be revised to include other facets of biodiversity^53^.

Our methods provide a rigorous, updatable approach for measuring both quantitative and qualitative progress toward meeting globally-informed area-based targets for nations, ecosystems, and species, while enabling local management decisions to reflect the world’s heterogeneous landscape of cultural and social needs, values, and knowledge. While our work was explicitly designed to inform and support actions needed to achieve both the CBD’s stated 2030 Agenda for Sustainable Development and their 2050 Vision for Biodiversity, the recommendations apply at any level of decision making, and the methods readily adapt to support more regionally focused policy. Indeed, the successful implementation of a global strategy will require multi-faceted conservation efforts from a variety of actors at every scale.

## Materials and Methods

### Species distributions

Our analyses included 32 649 terrestrial vertebrate species, comprising 6 436 amphibians and 6 169 mammals^39^, 9 987 birds^54^, and 10 057 reptiles^55^. We delineated broad native ranges based on expert maps, splitting the bird ranges by habitat use (breeding vs. nonbreeding) and treating these as separate species ranges, resulting in an effective total of 11 670 bird species. Expert maps were first translated to a coarse 55×55 km equal-area grid (0.5° grid cells at 30° latitudes) to reduce false presences^21^, and then further subdivided into planning units (PUs) of 27.5×27.5 km, resulting in 64 690 289 species-PU expert range (ER) combinations.

To represent fine-scale distributions of species, we extracted expert information on species-habitat associations for land cover^56^, tree cover^56^, and elevation^56,57^. We used land cover preferences to develop an association of each species with the 22 global land cover classes of the ESA CCI land cover product^58^. ESA CCI is an authoritative global land cover classification, providing information on dominant class for each terrestrial 300m pixel. For each land cover class we assessed relative suitability with each species habitat category in 25% intervals from 0 to 100% (Table S1). For example, mosaic natural/non-natural vegetation classes would get assigned partial suitability for multiple species habitat categories. We used species’ tree cover preferences to determine the minimum and maximum level of percent tree cover per 1 km^2^ (to the closest 10%) that would suffice as suitable habitat. Finally, we used species’ elevational preferences to determine suitable elevation habitat for each species. Following the rationale of Rondinini *et al*. (2005)^59^ and Jetz *et al*. (2007)^60^ we then determined the percent of habitat-suitable range (HSR) per 1 km^2^ pixel given these species elevation and habitat requirements ref 12. Specifically, we calculated average suitable area based on the combination of suitable elevation (based on the GMTED2010 dataset, as processed through Amatulli et al. (2018)^61^; see http://www.earthenv.org/topography), suitable land cover, and tree cover in 2019 (based on the ESA CCI landcover data and Landsat-based tree cover data^62^, respectively). We then aggregated these 1 km^2^ percent HSR values to the same 27.5×27.5 km PUs for further analysis, resulting in 51 554 595 species-PU HSR combinations. All species land-cover and elevation data and resulting HSR maps are accessible on Map of Life under the ‘Habitat Distribution’ panel, e.g. https://mol.org/species/range/Hybomys_planifrons.

### Protected areas and other effective area-based conservation measures

We used the World Database on Protected Areas to delineate current protected areas (PAs)^9^. Beginning with the June 2021 WDPA monthly release, we followed the WDPA’s recommendations on cleaning data for calculations of global coverage: we excluded PAs that did not have designated, inscribed, or established status, points without a reported area, marine reserves, and UNESCO Man and Biosphere Reserves. A buffer was created around PA point data with the area of the buffer equal to the reported area of the PA^63^. The PA polygons and buffered points were dissolved together, and intersected with GADM 3.0 coastline. The results were then rasterized to a 1×1km grid using a Behrmann equal-area projection.

We also used the World Database on other effective area-based conservation measures (WD-OECM). OECMs represent an important strategy for conservation, complementing PAs through active governance and management plans that result in sustained, positive conservation outcomes. Although WD-OECM only contains spatial data for five different countries as of June 2021, it is worthwhile to recognize their contribution on principle, and were so included here despite currently incomplete global coverage. OECM data were cleaned using the same approach as PAs.

### Human pressures

We used Kennedy *et al*. (2019)’s index of global human modification (HM) to characterize the cumulative impact of human pressures on the landscape^10^. We reprojected the map to the same 1×1km Behrmann equal-area projection used for the PAs using bilinear interpolation for HM values. We extended the map to cover the GADM 3.0 coastline, filling in any missing values (e.g., Antarctica) as NA, and masked out all current PAs. Values were then categorized into six levels of HM, following Kennedy *et al*.’s classification scheme: none (HM = 0), low (0 < HM ≤ 0.1), moderate (0.1 < HM ≤ 0.4), high (0.4 < HM ≤ 0.7), very high (0.7 < HM ≤ 1), and uncategorized (HM = NA). Uncategorized areas were generally typified by large freshwater lakes and ice.

### Economic data

We used 2019 estimates of gross national income adjusted for purchasing power parity (GNI PPP) expressed in current (2019) international dollars as a measure of the financial wealth of countries, as calculated by The World Bank (https://data.worldbank.org/indicator/NY.GNP.MKTP.PP.CD).

### Species representation targets

We used area-based representation targets to define the amount of habitat each species needed for protection to be considered safeguarded for the future. The amount of habitat potentially available for conservation was calculated by the count of 1-km HM pixels within each PU. Similarly, the amount of currently protected habitat was calculated by the count of 1-km PA pixels within each cell. Species within-cell areas was then calculated as the summed counts of PA and gHM pixels within cells, with total range area summed across all cells. The representation targets were then calculated as a function of a species’ total range size with a piecewise log-linear function that specified representation targets of 100% for species with ranges up to 10 000 km^2^ and 15% for species with ranges greater than 250 000 km^2^, chosen *a priori* to reflect the 15% of global surface area currently protected^9^; in theory, given a globally uniform distribution of PAs, widespread species would remain neutral to the analysis. Representation targets were capped at 1 000 000 km^2^, which prevented the optimization results from being overly influenced by the most widespread species. This choice affected 96 species, comprising 76 birds, 19 mammals, and 1 reptile.

### Optimization

We formulated the optimization problem as a linear program (LP)^8^ so that each of the *N* = 206 651 PUs was represented by a decision variable *x_i_*. The resulting LP was

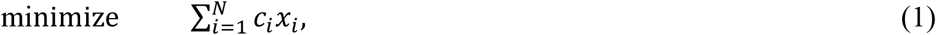

subject to:

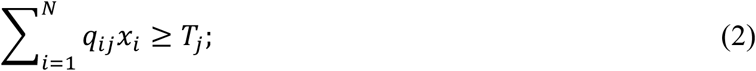

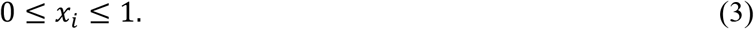

Equation (1) minimizes the area of the reserve network. The costs *c_i_* associated with protecting cell *i* reflected the amount of land in the cell that was not currently protected and hence was theoretically “available” for additional conservation action. These were computed with 1km-pixel counts of non-PA land for each cell.

The inequality in Eq. (2) ensures that the amount of area protected by the reserve network is above the representation target *T_j_* for each species *j*. The amount of habitat *q_ij_* available for future protection for species *j* in cell *i* and was calculated as

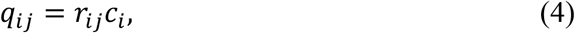

where *r_ij_* indicates the amount of habitat of species *j* in cell *i*. The species representation target *T_j_* is given by

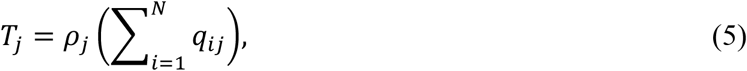

with *ρ_j_* the proportion of range area determined by the representation target.

Equation (3) constrains the decision variables to represent proportions between 0 and 1. Optimization was performed using Gurobi — a software package that employs the simplex and branch-and-bound algorithms for linear and mixed integer optimization^64^ — via the “prioritizr” R package^65^. All analysis was performed in R.

### Ecoregions and biomes

Achieving ecological representativeness in accordance with Aichi Biodiversity Target 11 is commonly interpreted as protecting at least 17% of each of the planet’s distinct ecoregions. To assess the comprehensiveness of biogeographical representation in the reserve network we calculated the amount of land needed in 846 ecoregions, using the Ecoregions2017 dataset^66,67^. We note that this dataset was not created with GADM 3.6 as the base world layer, which led to occasional small rounding errors in ecoregion areas in places of border discrepancies. This resulted in some ecoregions needing an area slightly greater than 100%; such regions were rounded down to 100%.

We found that only 60 ecoregions required protection of 17% or less of their total area, illustrating that the terrestrial target suggested by the expiring Aichi Biodiversity Target 11 is woefully inadequate. Grouping broadly by biome, none of the 14 distinct biome categories required less than 17% of total area.

### Conservation burden

We define the *conservation burden* of an administrative region as the ratio of the weighted sum of the region’s area contained in the reserve network and the region’s gross national income adjusted for purchasing power parity (*GNI*). This is calculated as

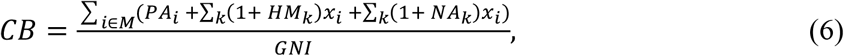

where *M* is the set of cells that overlap with the administrative region, *PA_i_* is the amount of currently protected area in cell *i*, *HM_k_* are the human modification pixel values in cell *i*, *NA_k_* are the pixels of terrestrial area in *i* with no HM values, and *x_i_* are the decision variables at optimality. Land with higher HM is weighted more heavily to reflect additional expenses associated with restoration of more heavily degraded habitat. The area with no HM values introduces a source of uncertainty in the estimate of conservation burden; for example, areas of rock and ice may be considerably less costly to conserve than freshwater areas, both of which are represented by NA values in the HM layer. As such, we calculated CB for two scenarios: one in which all *NA_k_* = 0 (for a lower bound on CB), and in which all *NA*_k_ = 1 (for a higher bound). These bound estimates are available in Table S2.

Conservation burden reflects the theoretical economic capacity of a country to achieve its national prescribed conservation objectives and is expressed in square meters of land per dollar. In practice, the economic capacity of a country will be heavily influenced by local land costs and the proportion of GNI that a country chooses to invest toward conservation, in addition to other factors. Summing across all global area needed and all GNI, the global conservation burden (GCB) provides a useful reference point for individual regions; GCB is 0.367 m^2^/$.

We also explored a formulation of conservation burden that excluded the contribution of current PAs to distinguish contributions already established from those not yet achieved. We called this the *future* conservation burden, which only considers costs associated with future additional conservation actions. *Present* burden (Fig. 5), by contrast, also considers the costs associated with maintaining land that is already protected. Countries with smaller present burdens generally had even smaller future burdens, sometimes by several orders of magnitude (Fig. S3). This suggests that most of the land requirements of countries with small present burdens come from areas that are already protected. Future burdens are also available in Table S2.

## Supporting information

Supplementary Table 1 - Land cover classes

Supplementary Table 2 - Countries

Supplementary Table 3 - Ecoregions

## Data availability

All datasets of species distributions, protected areas, and human modification used in this study are publicly available from their original sources. Detailed and interactive information on species distributions used in the analyses are also available at Map of Life (https://mol.org).

The global optimization results in Figure 1 (shapefile of 0.25° cells, with area needed and priority ranks of each cell) will be publicly shared in the ArcGIS Living Atlas of the World (https://livingatlas.arcgis.com/), and can presently be explored interactively on Map of Life and the Half-Earth Project’s mapviewer (https://www.half-earthproject.org/maps/).

The local optimization results in Figure 2 (raster of 1 km HM values within the reserve network) will be publicly shared in the ArcGIS Living Atlas of the World (https://livingatlas.arcgis.com/), and will be able to be explored interactively on the Half-Earth Project’s mapviewer (https://www.half-earthproject.org/maps/) by date of publication.

The ecoregion, biome, and country data in Figures 4 and 5 are shared as supplementary tables with this manuscript.

All other supporting data and code are available from the corresponding author upon request.

## Supplementary discussion

### Richness, endemism, and rarity

Species richness describes the number of species occupying a given region^26^, and is typically calculated by summing occurrences by grid cell. Because our probabilistic downscaling approach assigns a probability of occurrence to each 55km cell, summing occurrence probabilities by cell results in non-integer values of richness. Specifically, the species richness of cell *i* was calculated as

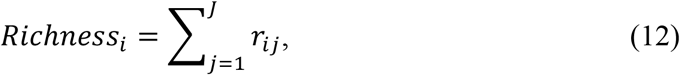

where *r_ij_* is the presence probability of species *j* in cell *i* and *J* is the total number of species (Fig. S1A).

Species endemism (also known as total range-size rarity or weighted endemism) describes the proportion of a species’ range that is found in a given region, summed across all species within the region^26^. The species endemism of cell *i* was calculated as

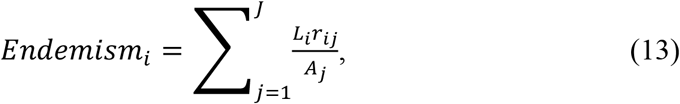

where *L_i_* is the area of land in cell *i* and *A_j_* is the total range area of species *j* (Fig. S1B). Species rarity (also known as average range-size rarity) is simply the endemism divided by the number of species present in each cell, and given by

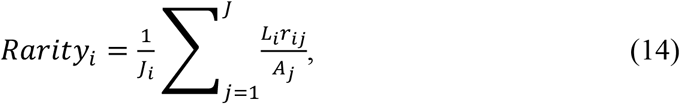

where *J_i_* is the number of species with a nonzero occurrence probability in cell *i* (Fig. S1C).

**Figure S1.**
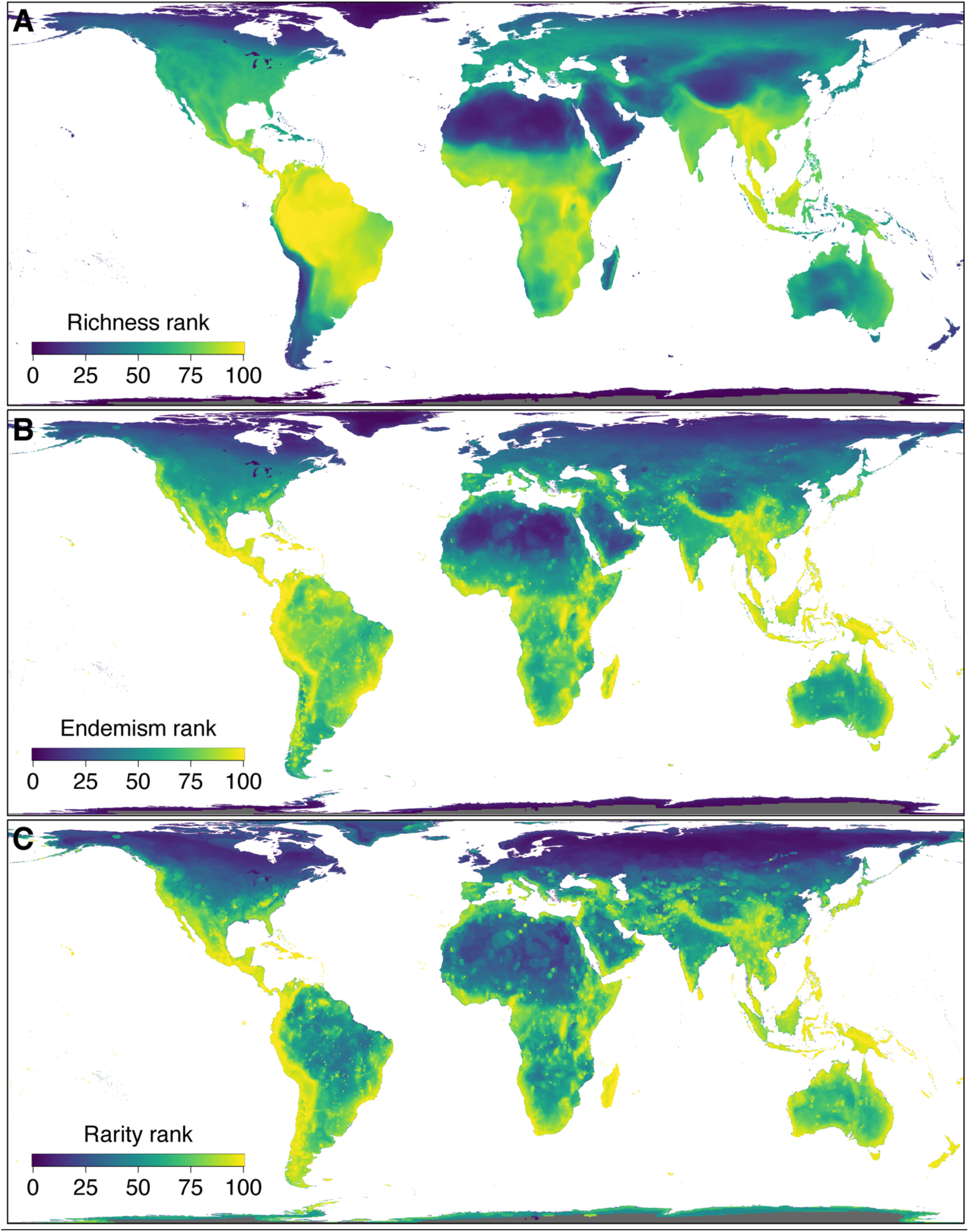
Percentile rank of species (A) richness, (B) endemism, and (C) rarity mapped for expert ranges of study species. Cells in grey did not contain any species occurrences, and were omitted from percentile rank calculations. Priority rank (Fig. 1A) most closely correlated to endemism rank (r_s_ = 0.846 for ER, 0.856 for HSR).

**Figure S2.**
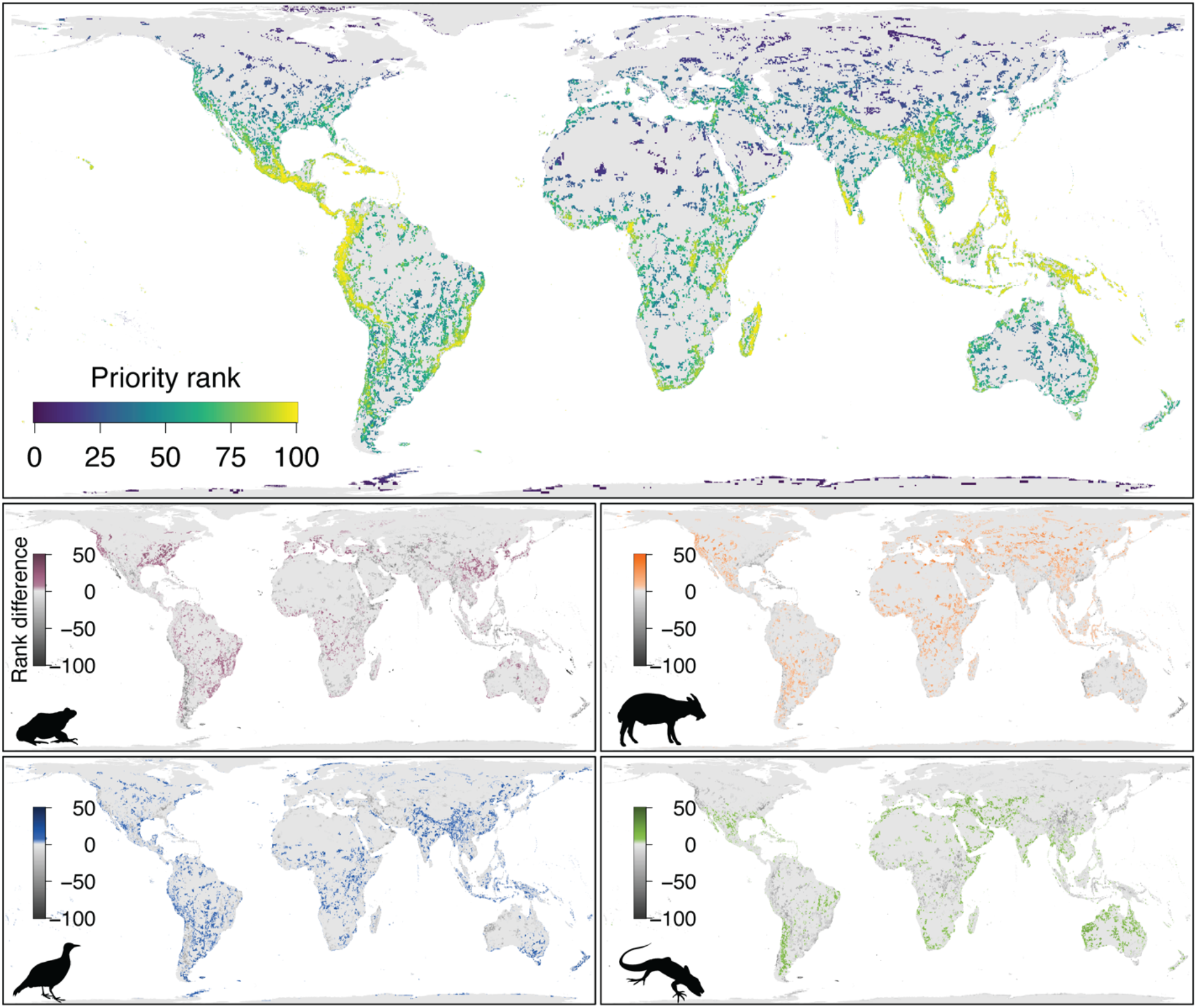
Differences in priority rankings between taxonomic groups, highlighting taxon-specific conservation hotspots. Such patterns can help guide local conservation strategies and support conservation advocacy for targeted groups of species. Rank difference of a species group is calculated as the priority ranks summed across all study species minus the priority ranks summed across species within that species group. Areas with rank difference > 0 indicate locations of greater conservation importance for a given species group relative to the rank of all vertebrates in that location.

**Figure S3.**
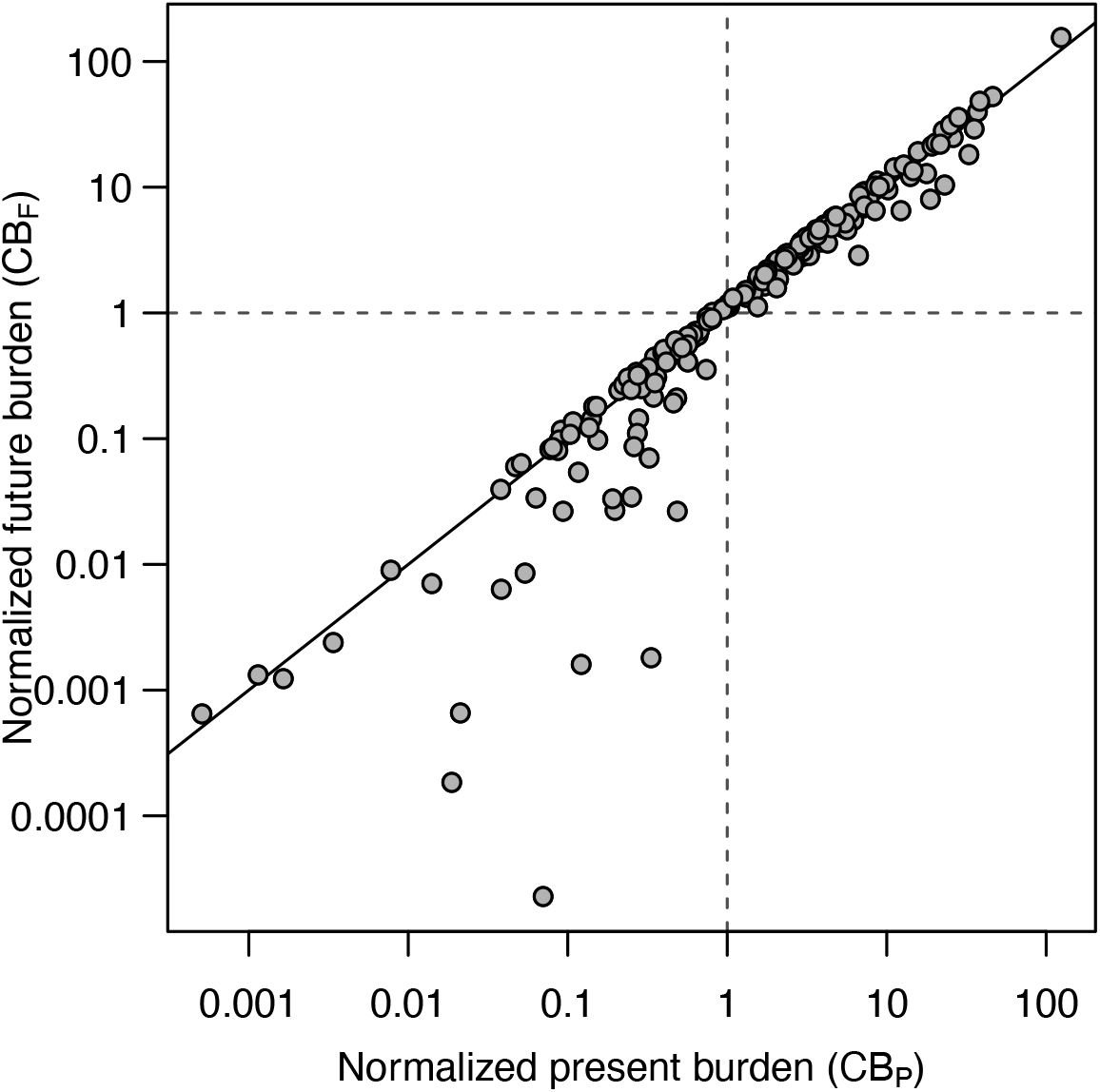
Present burden vs. future burden of countries, each normalized by present and future global conservation burdens GCB_P_ and GCB_F_, respectively. GCB_P_ and GCB_F_ are indicated by the dashed lines. Present burden accounts for both current PAs and additional areas needed; future burden reflects only additional areas needed.

## Acknowledgments

The authors would like to thank numerous employees at Map of Life for help with data curation and project support, in particular Michelle Duong and Jeremy Malczyk. We are grateful to contributors to recent workshops organized by the GEO BON Species Populations Working Group (at iDiv Leipzig), the National Geographic Society and Chinese Academy of Sciences (NGS-CAS, Beijing), CAS and Max Planck Society (Shanghai) that helped shape this research. This study benefited from NSF grants DBI-1262600 and DEB-1441737, EO Wilson Biodiversity Foundation grants, and NASA grants 80NSSC17K0282 and 80NSSC18K0435 to W.J.

## Author contributions

**D. Scott Rinnan:** Conceptualization, Data curation, Formal analysis, Investigation, Methodology, Project administration, Software, Validation, Visualization, Resources, Writing – original draft. **Yanina Sica:** Data curation, Formal analysis, Investigation, Methodology, Resources, Writing – review & editing. **Ajay Ranipeta:** Data curation, Formal analysis, Investigation, Methodology, Resources. **John Wilshire:** Data curation, Investigation, Methodology, Resources. **Walter Jetz:** Conceptualization, Data curation, Formal analysis, Funding acquisition, Investigation, Methodology, Project administration, Supervision, Visualization, Resources, Writing – review & editing.

